# regulaTER: An R Library to Study the Regulatory Roles of Transposable Elements

**DOI:** 10.1101/2024.06.29.601318

**Authors:** Doğa Eskier, Seray Yetkin, Nazmiye Arslan, Gökhan Karakülah, Hani Alotaibi

## Abstract

Gene expression is regulated at the transcriptional and translational levels and a plethora of epigenetic mechanisms. Regulation of gene expression by transposable elements is well documented. However, a comprehensive analysis of their regulatory roles is challenging due to the lack of dedicated approaches to define their contribution. Here, we present regulaTER, a new R library dedicated to deciphering the regulatory potential of transposable elements in given phenotype. regulaTER utilizes a variety of genomics data of any origin and combines gene expression level information to predict the regulatory roles of transposable elements. We further validated its capabilities using data generated from an epithelial-mesenchymal and mesenchymal-epithelial transition cellular model. regulaTER stands out as an essential asset for uncovering the impact of transposable elements on the regulation of gene expression, with high flexibility to perform a range of transposable element-focused analyses. Our results also provided insights on the contribution of the MIR and B element subfamilies in regulating EMT and MET through the FoxA transcription factor family. regulaTER is publicly available and can be downloaded from https://github.com/karakulahg/regulaTER.

## Introduction

Transcriptional regulation is a complex process underlying many biological events and one of the key factors of organismal complexity. By varying the expression levels of the genes in the genome, cells with identical genotypes can achieve a wide range of phenotypes, both during development and beyond. Besides transcription factors (TF) and many epigenetic factors, transposable elements (TEs) are involved in transcriptional regulation (Gebrie 2023). TEs are repeat genomic elements with the unique ability to reposition and create novel copies of themselves in the genome. Initially described by Barbara McClintock in 1956 (McClintock 1956), the impact of TEs on the genome and the cell at large has received more attention in recent years, revealing many of their roles in various biological pathways and processes. TE contributions to genomic complexity can occur via various mechanisms, including forming novel chromosomal features, novel genes, and regulatory elements (Liao et al. 2023). TEs are implicated in developmental processes, where they contribute to regulatory networks through transcription factor binding sites (TFBSs) for TFs involved in cellular differentiation (Fueyo et al. 2022, Gebrie 2023). In addition to their roles gained via exaptation, TEs can also influence cellular phenotype via transposition and novel insertion events, and aberrant TE activity can lead to genetic diseases or tumorigenesis (Bhat et al. 2022, Yandim and Karakulah 2019). Because of the potential deleterious effects of TE activity and insertion, these elements often must maintain a strict balance between activity and propagation or inhibition (Yoth et al. 2022). Therefore, most organisms have developed multiple mechanisms to restrict TE activity within their genomes, especially transposition and novel insertions, mainly KRAB-ZFPs, which methylate TE sequences in a targeted fashion, and PIWI-interacting RNAs, which silence TEs and prevent mobilization (Aravin et al. 2007, Molaro and Malik 2016, Yang et al. 2017).

As repeat genomic elements with high polymorphic variability, analysis of TEs often requires specialized tools and workflows (Goerner-Potvin and Bourque 2018, Lanciano and Cristofari 2020). Despite the range of tools available, various perspectives are missing in the current TE toolbox. For example, while the association of TE-derived TFBSs with gene regulatory networks is well-known, there are currently no tools to identify the contributions of TEs to transcriptomic regulation in a given phenotype. To overcome the current challenges, we developed regulaTER, an R library, to analyze multi-omics data to identify the potential regulatory impact of TEs. RegulaTER utilizes genomic peaks, such as those obtained from ATAC-seq or ChIP-seq libraries, and genes of interest, such as those identified by differential expression analysis of RNA-seq libraries, and accurately identifies TEs with potential role in gene regulation. Here, we describe and validate the functions of regulaTER using an epithelial-mesenchymal transition (EMT) and mesenchymal-epithelial transition (MET) experimental setting. We also provide novel insights into the contribution of TEs in the intricate regulation of EMT and MET.

## Materials and Methods

### The implementation of the regulaTER library

The regulaTER tool was created in the R environment (version 4.0.0). We used the biomartr package (version 1.0.2) (Drost and Paszkowski 2017) to import RepeatMasker annotations into the R environment for use in later calculations, GenomicRanges (version 1.42.0) (Lawrence et al. 2013) for processing of genomic peak ranges, and dplyr (version 1.1.1) for easier manipulation of genomic repeat category data (subfamily, family, type). For the calculation of TE enrichment, we used the binom.test and p.adjust functions available as part of the stats package of the base R installation. Genomic peaks were randomized for enrichment analyses using the BEDtools toolkit (version 2.29.0) (Quinlan and Hall 2010). For motif enrichment analyses, we used the HOMER genome analysis software (version 4.10.3) (Heinz et al. 2010) via marge (version 0.0.4.9999), an R environment API for HOMER. The library was documented, version-controlled, and uploaded to the GitHub software repository database via integrated Rstudio (version 2022.12.0) functions. The regulaTER library, as well as more detail on the use of its functions, are illustrated in Figure 1.

**Figure 1.**
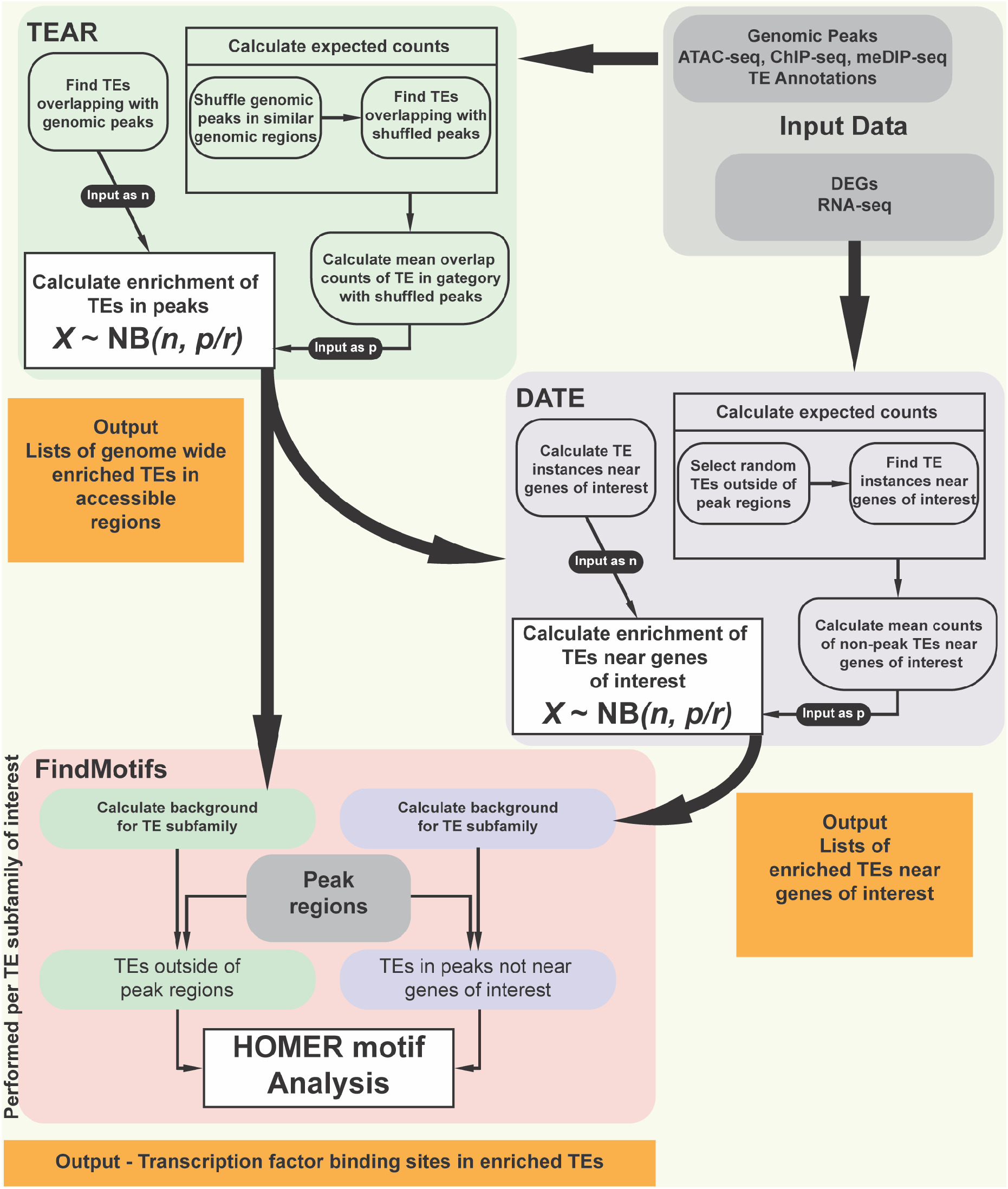
Detailed outline of the regulaTER functions and workflow. The gray boxes represent primary user-defined inputs (genomic peak data, DEG data). The green boxes indicate information regarding the TEAR function, while the purple boxes indicate information regarding the DATE function. The pink box represents the FindMotifs function. Orange boxes display the primary output of each function. Boxes with headers within the TEAR and DATE function descriptions each indicate a Monte Carlo simulation, which is repeated a number of times as determined by the user.

#### Identification of TEs Enriched in Accessible Genomic Regions via the Transposable Elements Enriched in Accessible Regions (TEAR) Function

The primary inputs of the TEAR function are the genomic sequencing peak files annotated by ChIPseeker (Yu et al. 2015), a list of files with genomic coordinates corresponding to ChIPseeker annotations and the sizes of each chromosome, and repeat annotations of the genome. In the function, TEs overlapping genomic peaks of interest showing differential accessibility are calculated via the Genomic Ranges package (“observed”). Afterward, the peak regions are randomized across the genome, with each peak being shuffled into a new location sharing the same genomic context, as compatible with the ChIPseeker annotations. Genomic region annotations for this randomization were obtained using the Table Browser utility of UCSC Genome Browser. TEs overlapping the randomized peak regions are then calculated. After the TE-randomized peak overlaps are calculated a number of times, determined by the numberOfShuffle input, the number of these overlaps is averaged (“expected”). To identify whether the “observed” number of TEs overlapping the peaks is significantly higher than the random distribution, we perform a negative binomial test (binom.test(observed, total TEs in genome, expected / total TEs in genome, alternative = “greater”). The p-values are corrected for multiple testing bias using the Benjamini-Hochberg correction method of the p.adjust function. We consider TEs with an FDR ≤ 0.05 and observed value ≥ 10 to be enriched in accessible regions (TEAR). The function performs the enrichment analyses for the TE subfamily, family, and type and outputs a list with three enrichment tables for each category. The TEAR function can accept peak files in narrowPeak, summits, or broadPeak formats generated by MACS2 and annotated by ChIPseeker (“format” parameter).

#### Identification of Accessible TEs Enriched Near Differentially Expressed Genes via the Differentially Expressed Gene Associated Accessible TEs (DATE) Function

The primary inputs of the DATE function are the TEAR function output, the repeat annotations and peaks used by TEAR, and a custom genomic coordinate file for the genes of interest. In this function, the number of accessible TEs in proximity or within the promoter regions of genes of interest is calculated for each TE subfamily via the Genomic Ranges package (“observed”). Following this calculation, a number of TEs of the same subfamily that do not lie in accessible genomic regions equal to the “observed” count are randomly selected, and these selected TEs are also overlapped with regions neighboring genes or promoter regions a number of times as determined by the user. The average number of such overlaps is calculated (“expected”). The enrichment for these gene-adjacent accessible TEs is calculated using the same negative binomial test parameters as in the TEAR function.

#### Identification of Genomic Motifs Contributing to Gene Regulatory Functions of Accessible TEs via the FindMotifs function

To identify genomic motifs that might contribute to the regulatory functions of identified TEs, we applied the FindMotifs function using the HOMER genomic sequence analysis tool via marge, an R environment API for HOMER. For the use of FindMotifs with TEAR function output, the background regions are calculated as TEs of the same subfamily outside accessible region peaks. For DATE function output, the background regions are calculated as accessible TEs outside of gene neighboring regions, i.e., with a distance to the gene region greater than the distance parameter (5000 by default).

### Cell culture

To validate the function and outputs of regulaTER, we used the NMuMG (Normal Murine Mammary Gland) epithelial cell line as an EMT and MET model. The cells were maintained in high glucose DMEM containing 10% fetal bovine serum, 100U/ml penicillin, 0.1 mg/ml streptomycin, and 1% non-essential amino acids mixture in an incubator set to 37°C and 10% CO2. As a supplement, 10 μg/ml insulin (Sigma-Aldrich) was freshly added to the medium. EMT was induced by adding 2.5 ng/ml TGFβ3 (PeproTech) and progressed for 10 days. MET was induced by the removal of TGFβ3 from the medium. MET was observed for 4 days.

### Gene expression analysis

#### qPCR

RNA was prepared using the Nucleospin RNA II kit (Macherey-Nagel) as recommended by the manufacturer. One microgram of total RNA was used for cDNA synthesis using the Maxima First Strand cDNA Synthesis kit (Thermo Scientific). Primers and probes used for qPCR were previously reported (Alotaibi et al. 2015). Quantitative RT-PCR was performed using the AMPIGENE qPCR probe mix (Enzo), and amplification was performed in the 7500 Real-time PCR system (Applied Biosystems) using the following cycling parameters: 95°C 5min, 40 cycles of 95°C 15s, 60°C 60s. Relative expression to the vehicle control (normalized to Gapdh expression) was calculated using the ΔΔCt method. Data represents at least three biological replicates.

#### Western Blot

NMuMG cells for each time point were harvested and lysed in RIPA buffer. Protein concentration was determined by the BCA Protein Assay Kit (Takara). Protein samples were then diluted with Laemmli Sample buffer. After electrophoresis, proteins were transferred to a nitrocellulose membrane, blocked for 1 h in blotto (Tris-buffered saline containing 0.5% Tween 20 and 5% nonfat milk powder), and incubated with anti-E-cad antibodies (BD Biosciences, 250 ng/ml) overnight at 4 °C with shaking. Anti-GAPDH (Santa Cruz, 50 ng/ml) was used as a loading control. Detection was done by incubation with horse-radish peroxidase-conjugated secondary antibodies (1:10000, Dianova), ECL prime reagent (GE Healthcare), and exposure to X-ray films.

#### Confocal Imaging

Cells were washed with PBS and fixed with 4% formaldehyde (Sigma) for 10 min at room temperature, washed twice with PBS, and permeabilized with 0.25% Triton X-100 (Sigma) for 5 min. After washing twice with PBS, cells were incubated with primary antibodies in PBS with 5% BSA (1:200) for one hour at room temperature. Afterward, Alexa594-conjugated secondary antibody was applied for an additional one hour after washing in PBS. Nuclei were visualized with DAPI (1:1000, Invitrogen) and mounted with Prolong Diamond Antifade Mountant solution (Thermo Scientific). Confocal microscopy was carried out with an LSM 880 microscope equipped with ZEN software (Zeiss). The antibodies used were anti-E-cadherin (BD Bioscience), Alexa Fluor 488 Phalloidin (Invitrogen), DAPI (Invitrogen), and Alexa Fluor 594 goat anti-mouse IgG (Invitrogen)

### ATAC-seq

ATAC-seq reactions and library preparation were performed using the ATAC-sequencing Kit (Active Motif), following the manufacturer’s protocols. Briefly, the nuclei of approximately 100000 cells were isolated and treated with Tn5 transposase, followed by adaptor ligation and PCR amplification of the libraries. The prepared libraries’ quality control was checked using the Agilent Bioanalyzer 2100, and the libraries were sequenced using the Illumina NovaSeq 6000 sequencing platform (acquired by a third-party service provider). We performed the alignment and peak calling analysis of raw ATAC-seq reads using the nf-core atacseq (version 2.1.0) workflow, using the command “nextflow run nf-core/atacseq --input design.csv --genome mm10 -profile docker --save_reference”. During the workflow, the reads were aligned to the UCSC Genome Browser mm10 mouse genome assembly using the BWA aligner, and the peaks were called in “broad” format using the MACS2 peak calling software.

### RNA-seq

Isolation of RNA samples was performed with the Nucleospin RNA II kit (Macherey-Nagel) as recommended by the manufacturer. RNA-seq libraries were prepared using the Illumina Stranded Total RNA Prep with Ribo-Zero Plus kit following the manufacturer’s protocols. Quality control of the prepared libraries was checked using the Agilent Bioanalyzer 2100, and the libraries were sequenced using the Illumina NovaSeq 6000 sequencing platform (acquired by a third-party service provider). The RNA-seq libraries were trimmed of low-quality bases and reads and possible adapter contamination using Trimmomatic (version 0.39) (Bolger et al. 2014). The remaining high-quality reads were aligned to the GENCODE GRCm38 (version M21) mouse genome assembly obtained from the GENCODE database, using HISAT2 (version 2.0.5) (Kim et al. 2015), using the “hisat2-build” command to index the genome, and “hisat2 -x -U --dta -S -p 6” to align the reads. The resulting SAM alignment files were converted to binary BAM format using SAMtools utilities (version 1.10) (Danecek et al. 2021) “samtools view” and “samtools sort”. The RNA-seq libraries were then quantified using the featureCounts function of the Rsubread R library (Liao et al. 2019). Transcripts with FPKM values lower than 0.5 in both replicates of all samples were filtered out. We analyzed differential expression using the edgeR library (version 3.32.0) (Robinson et al. 2010) using the count data of remaining transcripts. The count data was normalized using the TMM normalization method and then fit for the generalized linear model (GLM). We considered genes with abs(log2FC) ≥ 1 and adjusted p-value (false discovery rate, FDR) ≤ 0.05 to be differentially expressed genes (DEG). The biological implications of the DEGs were summarized using Gene Set Enrichment Analysis (GSEA) with the GSEA function of the clusterProfiler package (version 3.18.1) (Yu et al. 2012) on the hallmark genes of Mus musculus, with the line “GSEA(geneList = <named log2FC values>, minGSSize = 15, maxGSSize = 500, pvalueCutoff = 0.05, eps = 0, seed = TRUE, pAdjustMethod = “BH”, TERM2GENE = dplyr::select(mm_hallmark_sets, gs_name, gene_symbol))”.

### GO enrichment analysis

GO term enrichment analysis for TE-associated genes was performed using the enrichGO function of the clusterProfiler library (enrichGO(gene.list, OrgDb = org.Mm.eg.db, “SYMBOL”, “BP”)) and filtered for level 5 terms using the gofilter function of the same library (gofilter(enrichGO.results, level = 5)). For each TE analyzed, the associated genes were determined using the findOverlaps function of the GenomicRanges library to first identify accessible TEs in the subfamily of interest by finding TE instances overlapping ATAC-seq peaks showing differential accessibility in the sample, then by finding accessible TEs within 5000 nucleotides of upregulated genes (findOverlaps(maxgap = 5000, ignore.strand = TRUE)). The results were visualized using the circle_dat and GOBubble functions of the GOplot library (version 1.0.2) (Walter et al. 2015).

### Statistical analyses and visualization

Statistical analysis for gene expression analysis was performed using one-way ANOVA and Tukey’s multiple comparison tests. For data generated from computational studies, the R statistical computational environment is used for most statistical analyses and visualization purposes. We used pheatmap (version 1.0.12) to visualize the expression z-scores of genes during cellular differentiation using row-wise scaling. Summarized data was visualized using the ggplot2 (version 3.3.2) library.

UCSC Genome Browser images were generated using BigWig files representing each sample; files were uploaded to the UCSC genome browser (GRCm38/mm10), and tracks were visualized using the group-scaled and mean windowing function options. The ReMap track for each transcription factor was added using the same visualization parameters. Sample names and gene IDs were retyped in Adobe Illustrator for clarity and better resolution of the fonts.

## 3. Results

### Functions of regulaTER

To overcome the limitations while studying the contribution of TEs in gene regulation (stated in the introduction), we developed regulaTER, an R library for the analysis of multi-omics data to identify the potential regulatory impact of TEs. regulaTER utilizes organism-independent genomic peaks generated from a wide range of experiments, such as ChIP-seq, ATAC-seq, CUT&RUN, and meDIP-seq. Combined with gene expression data from RNA-seq, regulaTER delineates the contribution of TEs in gene regulation. The library has three core functions: Transposable Elements Enriched in Accessible Regions (TEAR), Differentially Expressed Gene Associated Accessible TEs (DATE), and FindMotifs (Figure 1). The TEAR function primarily determines the enrichment of TEs in genomic peaks, while the DATE function investigates the association of the TEs identified by TEAR with differentially expressed genes. The FindMotifs function calculates or predicts any potential regulatory motifs enriched in a given TE instance and can follow either the TEAR or the DATE function. These functions have been optimized to identify the enrichment of accessible TEs and the transcription factor binding motifs in a given phenotype. An overview of the functions is presented in Figure 1.

### Establishment and validation of EMT and MET cell model to test regulaTER functions

To showcase the functionality of regulaTER, we generated and used omics data representing EMT and MET, which are two biological processes with critical roles during embryonic development, wound healing, and tumorigenesis (Yao et al. 2024). The advantage of using this model is twofold. First, the transcriptional regulation of the EMT program is well documented, with multiple TEs implicated in its regulation (Apostolou et al. 2015, Reyes-Reyes et al. 2017, Simo-Riudalbas et al. 2022), making it a powerful validator. Second, while EMT and MET are two closely related processes, the regulation of MET is distinct from that of EMT and is poorly understood, with no TEs known to be involved in its regulation, thus rendering MET a powerful model for discovering TEs involved in gene regulation (Pei et al. 2019, Yao et al. 2011).

We used the normal murine mouse mammary gland cell line NMuMG to generate expression and chromatin accessibility profiles. The cells were treated with TGFβ3 for 10 days (hereafter called MNM10; MNM as in Mesenchymal NMuMG), and then TGFβ was removed from the medium by three washes with PBS; the cells at this point initiate MET and are incubated in regular culture medium to recover for 4 days (designated PT100; PT as in Post-Treatment).

Collected samples from control, MNM10, and PT100 cells were used for RNA and protein isolation to validate gene expression changes conferring the quality of EMT and MET. By examining the expression levels of Cdh1 and Cdh2, we confirmed the cadherin switch, a bona fide hallmark of EMT and MET (Figure 2A). Protein levels of E-cadherin showed similar changes to the RNA levels of Cdh1 (Figure 2B). In addition, confocal images revealed morphological changes associated with rearranged actin into stress fibers, validating the characteristic features expected during EMT and MET (Figure 2C).

**Figure 2.**
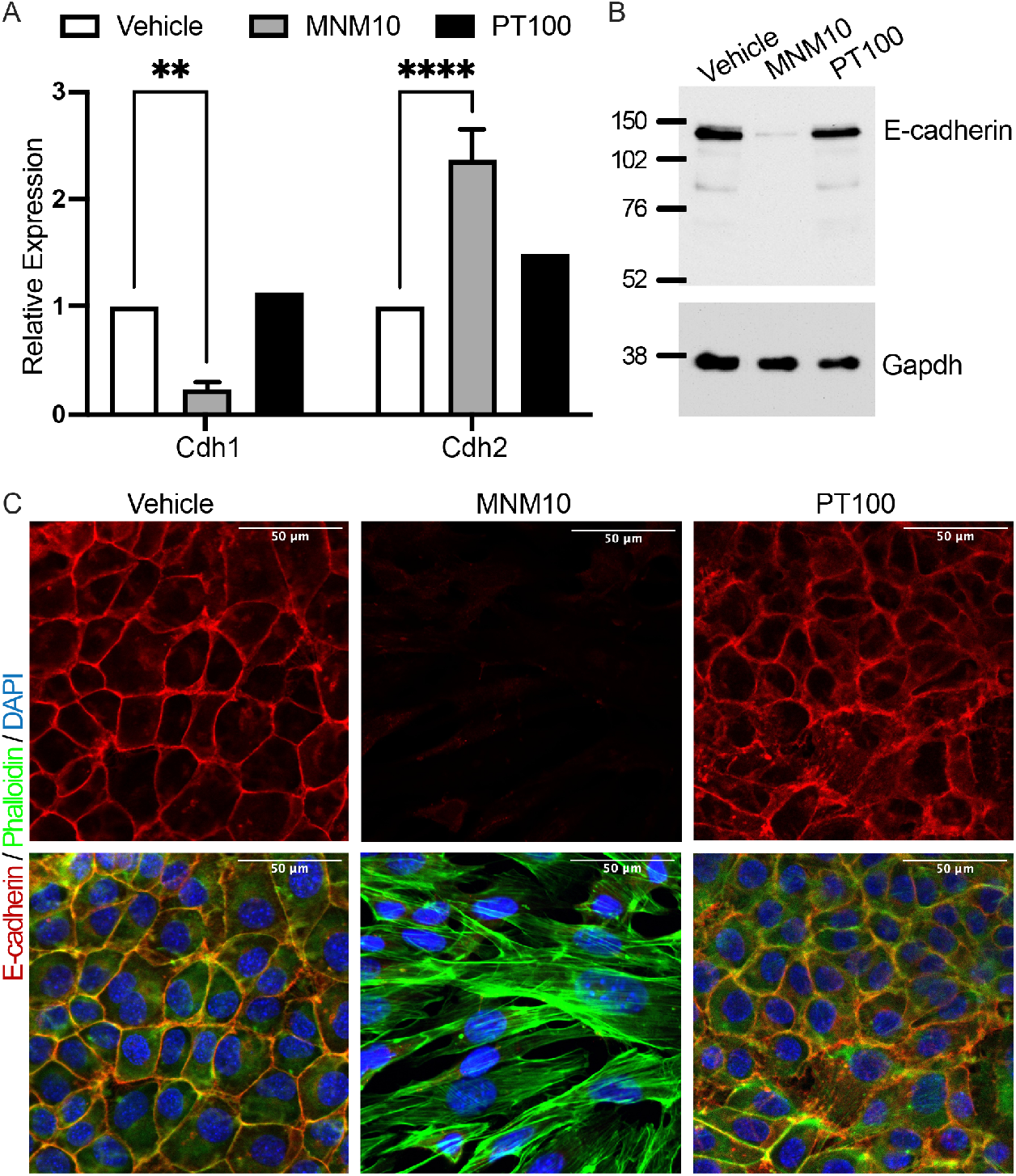
NMuMG cells successfully exhibited EMT and MET in response to TGFβ. A) qPCR analysis of the cadherin switch during EMT shows decreased Cdh1 and increased Cdh2 expression, which is reversed during MET. B) Changes in RNA levels were also reflected at the protein level, showing a decrease of E-cadherin expression during EMT, which is restored during MET. C) Confocal images of NMuMG cells showing the expression and localization of E-cadherin and Actin. Actin was visualized with Phalloidin, and the nuclei were stained with DAPI. The scale bar is 50 μm.

Collected and validated RNA from each sample was then used for RNA sequencing. We identified 3721 differentially expressed genes during EMT (2070 upregulated and 1651 downregulated, abs(log2FC) ≥ 1, FDR ≤ 0.05). On the other hand, 5133 genes were differentially expressed during MET (2929 upregulated and 2204 downregulated, abs(log2FC) ≥ 1, FDR ≤ 0.05) (Tables S1 and S2). The changes in the expression profiles corresponded to a typical EMT and MET (Figure 3A and 3B). GSEA confirmed that the differentially expressed genes correlated perfectly with the EMT hallmark gene set (Figure 3C and 3D).

**Figure 3.**
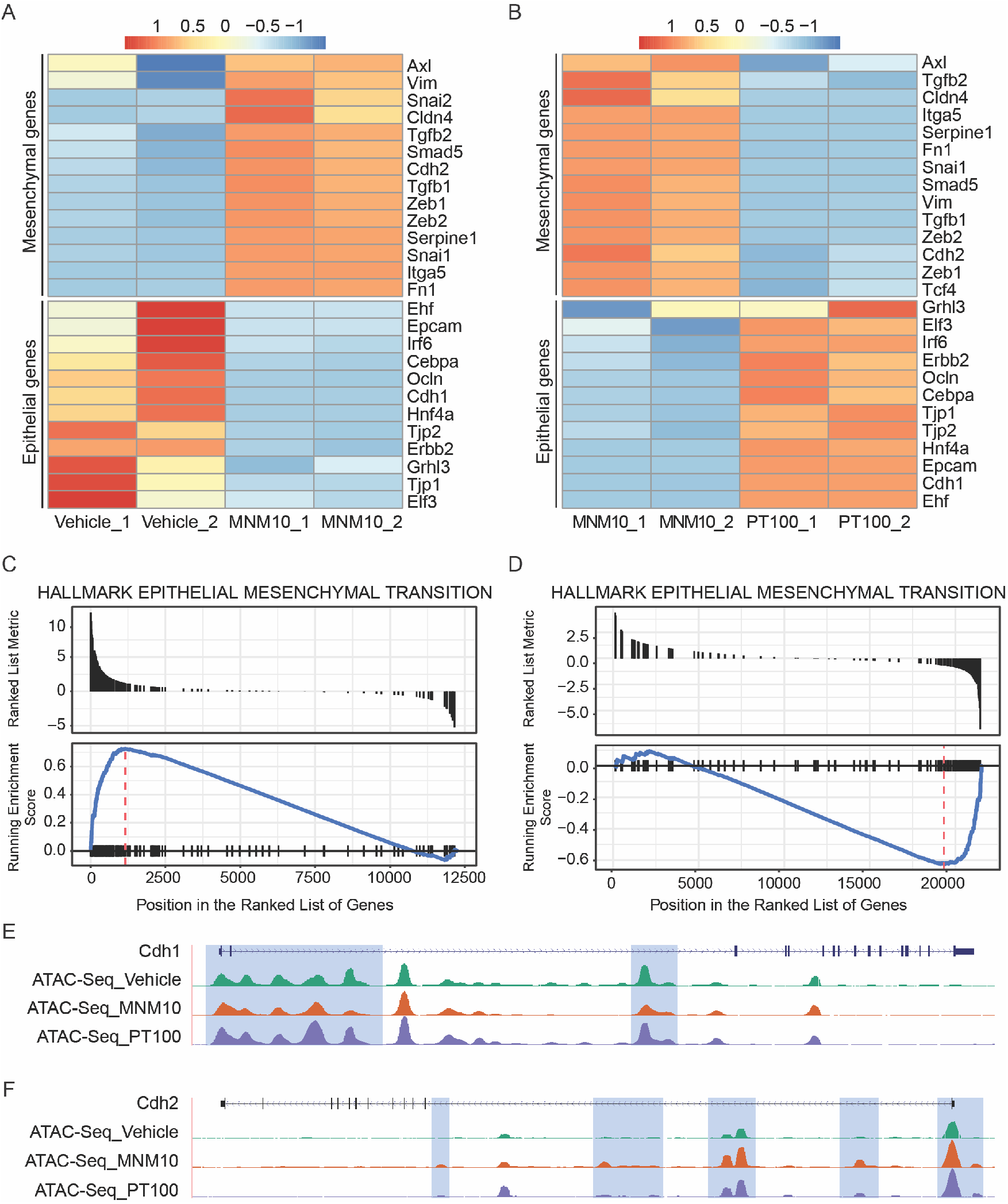
Gene expression analysis and differential chromatin accessibility during EMT and MET. Scaled expression heatmap of EMT markers in the (A) MNM10/Vehicle and (B) PT100/MNM10 comparisons. The colors of the cells indicate the z-score of the gene across the samples. GSEA analysis on the gene set of EMT hallmarks for Mus musculus in the (C) MNM10/Vehicle and (D) PT100/MNM10 comparison. The X-axis indicates the genes as ordered by the GSEA ranking list. The blue line in the bottom panel reflects the enrichment score of each gene. (E and F) UCSG genome browser images showing differential chromatin accessibility for Cdh1 (E) and Cdh2 (F).

We then prepared ATAC-seq libraries using cells from the same culture plates used for RNA-seq. The changes in gene expression paralleled with the changes in chromatin accessibility as well. For example, the Cdh1 promoter and intronic enhancers showed marked differential accessibility during EMT and MET (Figure 3E). Similarly, the promoter of Cdh2 was more accessible during EMT (Figure 3F). Examining the ATAC-seq data revealed 23636 differentially accessible peaks during EMT and 12129 during MET; 1.2% of these peaks were associated with genes known to be involved in EMT and MET (distance to gene body ≤ 5000 bp).

### MIR is the most significantly enriched TE family during EMT and MET

The generated ATAC-seq and RNA-seq data were used as inputs for regulaTER, and following the procedure summarised in the methods section (and detailed in the user manual), we implemented the different functions of regulaTER as outlined in Figure 1. To test and verify the TEAR function, we calculated differential peak accessibility of the ATAC-seq data by comparing MNM10 to Vehicle and PT100 to MNM10 using the count data of consensus peaks as input for the exactTest function of edgeR. Peaks with log2FC ≥ 1 and FDR ≤ 0.05 were considered to have increased accessibility. These peaks were annotated using the ChIPseeker package, which was then used as input for the TEAR function. The peaks were shuffled 10 times across the genome to calculate the expected value, as described in the methods section. As a result, 121 TE subfamilies were found to be enriched in ATAC-seq peaks in the MNM10 sample, while 45 subfamilies were enriched in the PT100 sample; 24 subfamilies were common in both samples (Tables S3 and S4). We then examined the top 10 TE subfamilies and families enriched in accessible peaks to identify whether any specific TEs had a potentially significant contribution to the EMT and MET processes. Out of the top 10 subfamilies in EMT and MET, five were common in both processes (RMER6C, MIRb, RMER15, MIR, and RLTR48A) (Table 1 and 2). Additionally, the MIR family was the most enriched in EMT and the second most enriched in MET. Among the accessible MIR TEs in EMT, three subfamilies (MIR, MIRb, MIR3) were within 5000 bp of 1423 genes, including Zeb1, a known EMT regulator. Meanwhile, the accessible MIR and MIRb instances in MET were found in the proximity of 343 genes, including Cdh1 (Table S5).

**Table 1.** Top 10 TE subfamilies enriched in accessible peaks in EMT.

| Repeat subfamily | Total genomic count | Observed | Expected | FDR | Fold change |
| --- | --- | --- | --- | --- | --- |
| RMER6C | 4930 | 176 | 35 | 5.59E-64 | 5.05 |
| MIRb | 41669 | 1051 | 636 | 6.43E-51 | 1.65 |
| RMER15 | 18012 | 466 | 237 | 6.49E-39 | 1.97 |
| MIR | 38078 | 865 | 571 | 1.01E-29 | 1.51 |
| RLTR17 | 1752 | 91 | 22 | 1.75E-27 | 4.22 |
| ORR1F | 22992 | 488 | 286 | 9.40E-27 | 1.71 |
| RLTR48A | 621 | 57 | 10 | 3.77E-24 | 5.87 |
| MLT1H | 2337 | 94 | 28 | 6.42E-22 | 3.31 |
| MIR3 | 15244 | 430 | 264 | 3.21E-20 | 1.63 |
| RLTR30D2_MM | 733 | 50 | 10 | 1.01E-18 | 5.22 |

**Table 2.**
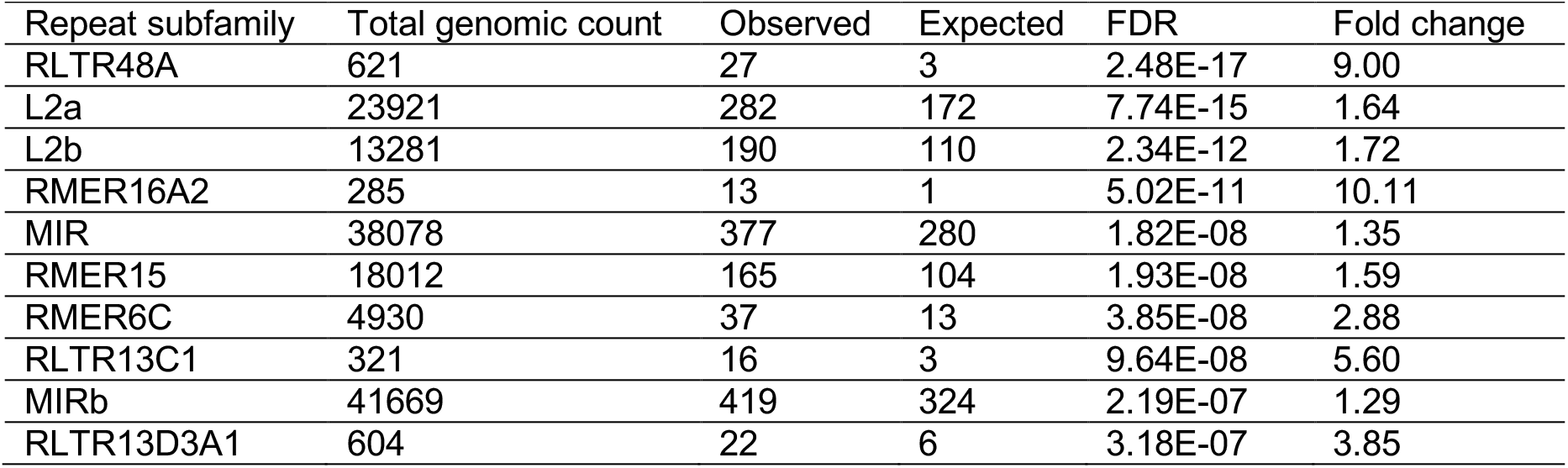
Top 10 TE subfamilies enriched in accessible peaks in MET.

| Repeat subfamily | Total genomic count | Observed | Expected | FDR | Fold change |
| --- | --- | --- | --- | --- | --- |
| RLTR48A | 621 | 27 | 3 | 2.48E-17 | 9.00 |
| L2a | 23921 | 282 | 172 | 7.74E-15 | 1.64 |
| L2b | 13281 | 190 | 110 | 2.34E-12 | 1.72 |
| RMER16A2 | 285 | 13 | 1 | 5.02E-11 | 10.11 |
| MIR | 38078 | 377 | 280 | 1.82E-08 | 1.35 |
| RMER15 | 18012 | 165 | 104 | 1.93E-08 | 1.59 |
| RMER6C | 4930 | 37 | 13 | 3.85E-08 | 2.88 |
| RLTR13C1 | 321 | 16 | 3 | 9.64E-08 | 5.60 |
| MIRb | 41669 | 419 | 324 | 2.19E-07 | 1.29 |
| RLTR13D3A1 | 604 | 22 | 6 | 3.18E-07 | 3.85 |

### B elements are enriched near DEGs in EMT and MET

Following the identification of TEs enriched in genome-wide accessible regions, we examined accessible TE enrichment near DEGs. As DEGs are an important indicator of a given phenotype, we wanted to identify whether accessible TEs were enriched near genes of interest differently than those enriched across the genome. To do so, we used the DATE function, which uses a different enrichment method to accurately identify DEG-associated TEs.

We applied the DATE function using the output of the TEAR to identify accessible TEs enriched in the neighboring regions of DEGs that show upregulation in either MNM10 or PT100 samples (regionSelection = “gene”, distance = 5000). We identified 32 TE subfamilies with enrichment of accessible instances near the DEGs in both MNM10 and PT100 (Tables S6 and S7). We selected the 10 TE subfamilies showing the highest enrichment in both samples for further analysis. Of these 10 subfamilies, 5 were commonly enriched in both conditions, namely B3A, B1_Mus1, ID_B1, B4A, and B3 (Figure 4A). To identify whether the genes showing potential regulation by these TEs significantly impacted the EMT (or the MET) transcriptome, we performed GO enrichment analysis using the upregulated genes neighboring the accessible instances of these TEs. Our results revealed biological pathways associated with EMT and MET (Figure 4B-C and Figure S1). Notably, response to TGFβ, regulation of substrate adhesion, migration, and wound healing are among the most enriched biological processes identified in EMT. Upregulated genes (such as Acta2, Itgb5, and Timp1) relevant to these processes also showed marked chromatin accessibility (Figure S2). In contrast, MET-related enrichment processes were linked to developmental processes or proliferation (Table S8).

**Figure 4.**
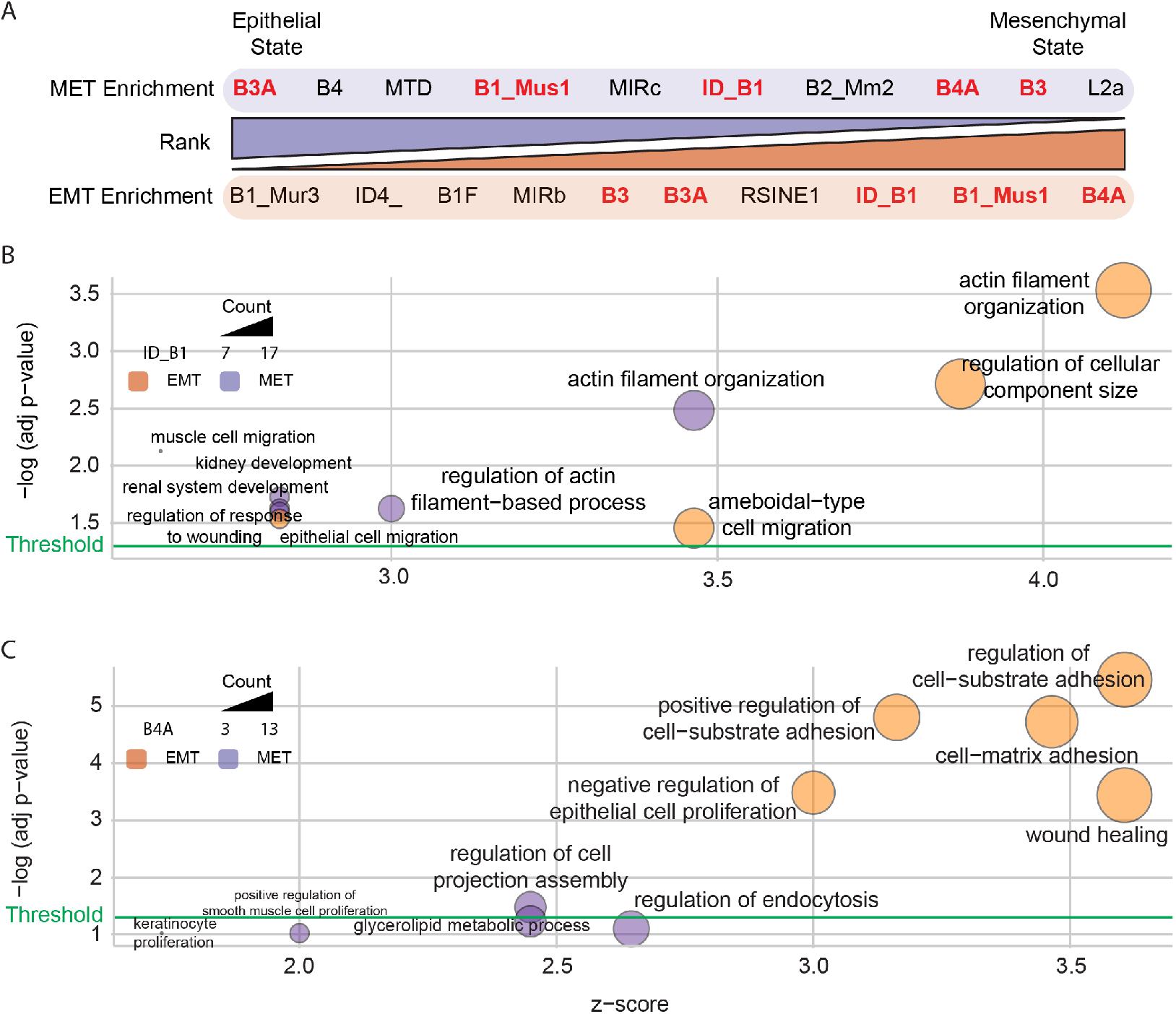
Five commonly enriched TEs show relevance to EMT- and MET-related biological processes. (A) The top 10 TE subfamilies enriched after the DATE analysis in EMT (orange) and MET (purple). TE subfamilies are ranked according to their contribution to epithelial (left end of the bar) or mesenchymal (right end of the bar) states. The TE subfamilies in red and bold text represent those commonly enriched in EMT and MET. (B-C) Modified GO bubble plots of the GO enrichment results for the upregulated genes within 5000 bp of the accessible instances of the B element TE subfamilies ID_B1 (B) and B4A (C). The x-axis indicates the z-score of the term, which indicates whether the genes assigned to the term are more upregulated (z-score > 0) or downregulated (z < 0). The y-axis represents the negative log of the adjusted p-value of the term, with the horizontal green line corresponding to an adjusted p-value of 0.05. The color of the bubbles indicates the process the term is associated with, with orange shapes enriched in EMT and purple shapes enriched in MET. The shape’s size indicates the number of TE-neighboring upregulated genes in the term.

### FoxA1 and FoxA2 confer EMT and MET regulation through TE-contributed binding sites

In order to predict the potential roles of the identified TEs in gene regulation, we applied the FindMotifs function using the output of the DATE function. The top 10 TEs enriched in EMT and MET were analyzed separately. We examined the FindMotifs results for the five TEs enriched near DEG clusters with significant contributions to EMT or MET. Our examination revealed that the accessible B1_Mus1 and B4A instances near the EMT DEGs showed enrichment for FoxA1 and TEAD binding sites. To further validate the contribution of these TE-provided motifs to gene regulation, we visualized the upregulated EMT DEGs alongside ATAC-seq peaks and ChIP-seq peaks (ReMap track) in the UCSC Genome Browser. FoxA1 enrichment was observed within an accessible TE in the first intron of Tgfb2 (Figure 5A), while TEAD enrichment was localized to the promoter and 2 introns of Cdh2 (Figure 5B). Examining the motifs within MET-enriched TEs, binding sites for FoxA2 stand out among other transcription factor motifs. FoxA2 motifs were identified in accessible ID_B1 located in the promoter and the first intron of Agt (Figure 5C) and in B4A located in the intron of Rab17 (Figure 5D).

**Figure 5.**
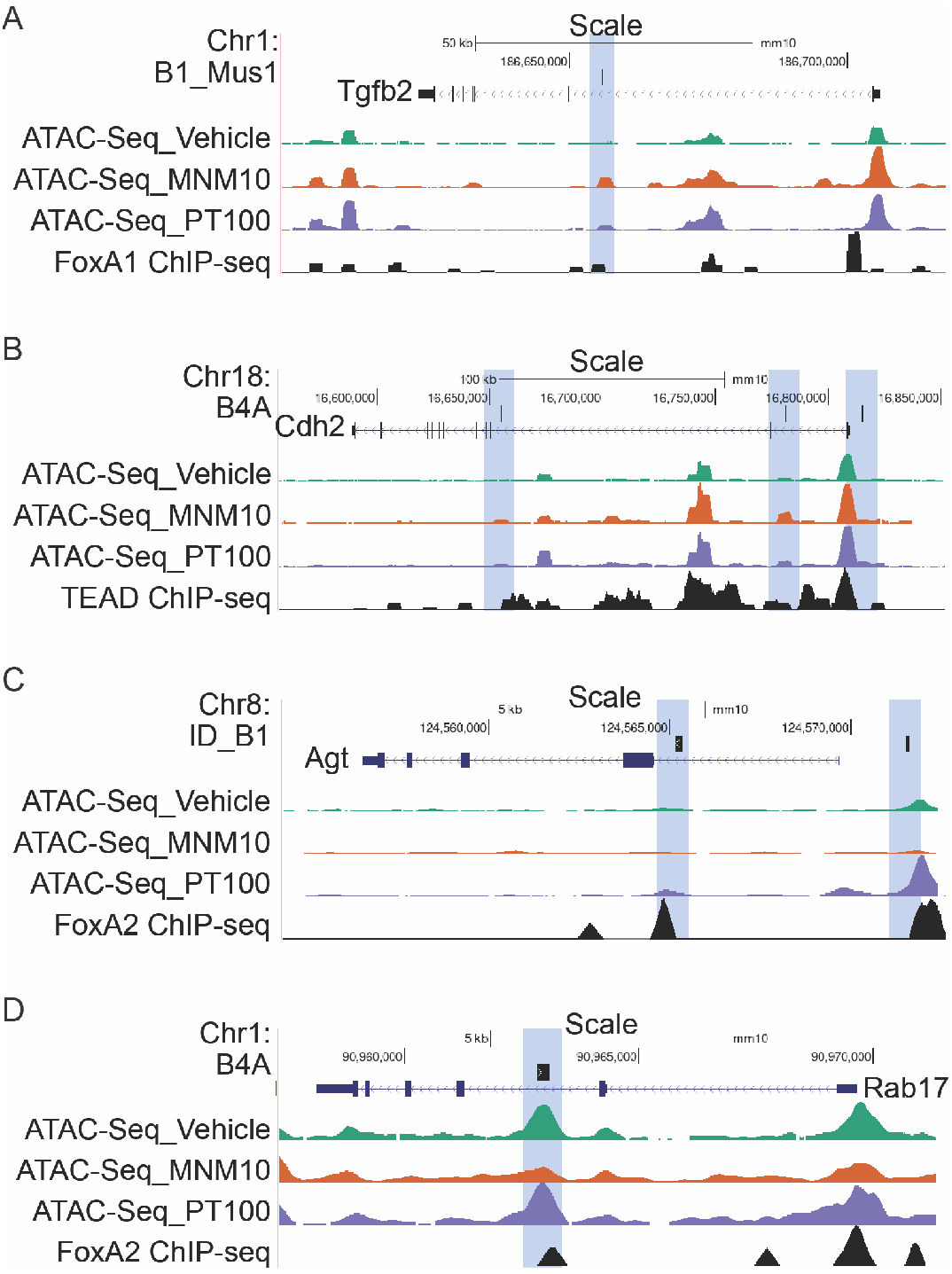
Regulation of EMT and MET by TEAD and the FoxA family of transcription factors through TE-contributed binding sites. UCSC Genome Browser images of (A) Tgfb2 and B1_Mus1, (B) Cdh2 and B4A, (C) Agt and ID_B1, and (D) Rab17 and B4A. Regions highlighted in blue indicate TE instances showing increased differential accessibility in EMT (A-B) and MET (C-D). The green tracks below the gene regions indicate ATAC-seq accessibility scores for the Vehicle sample, the orange tracks indicate accessibility for the MNM10 sample, and the purple tracks indicate the same for the PT100 sample. The black track indicates the ReMap ChIP-seq binding profiles for (A) FoxA1, (B) TEAD, and (C-D) FoxA2 TFs.

### Comparison to similar analysis methods

To highlight the uses and advantages of regulaTER, we compared it to similar analysis tools and databases. We selected the three available tools with a high amount of overlap with our library used to calculate the potential effects of TEs on the regulation of nearby genes, namely GREAM (Chandrashekar et al. 2015), TEffectR (Karakulah et al. 2019), and PlanTEnrichment (Karakulah and Suner 2017). Due to the different input types and computational methods utilized by each tool, a quantitative comparison was unfeasible. Therefore, we compared each tool’s capabilities and potential uses (Table 3). Overall, regulaTER has several unique strengths that were not found in other tools. Of highest relevance is the ability of regulaTER to accept both custom genomes and repeat annotations and its capacity to accept genomic sequencing peaks to consider the epigenomic state of individual TE loci in the enrichment analysis.

**Table 3.**
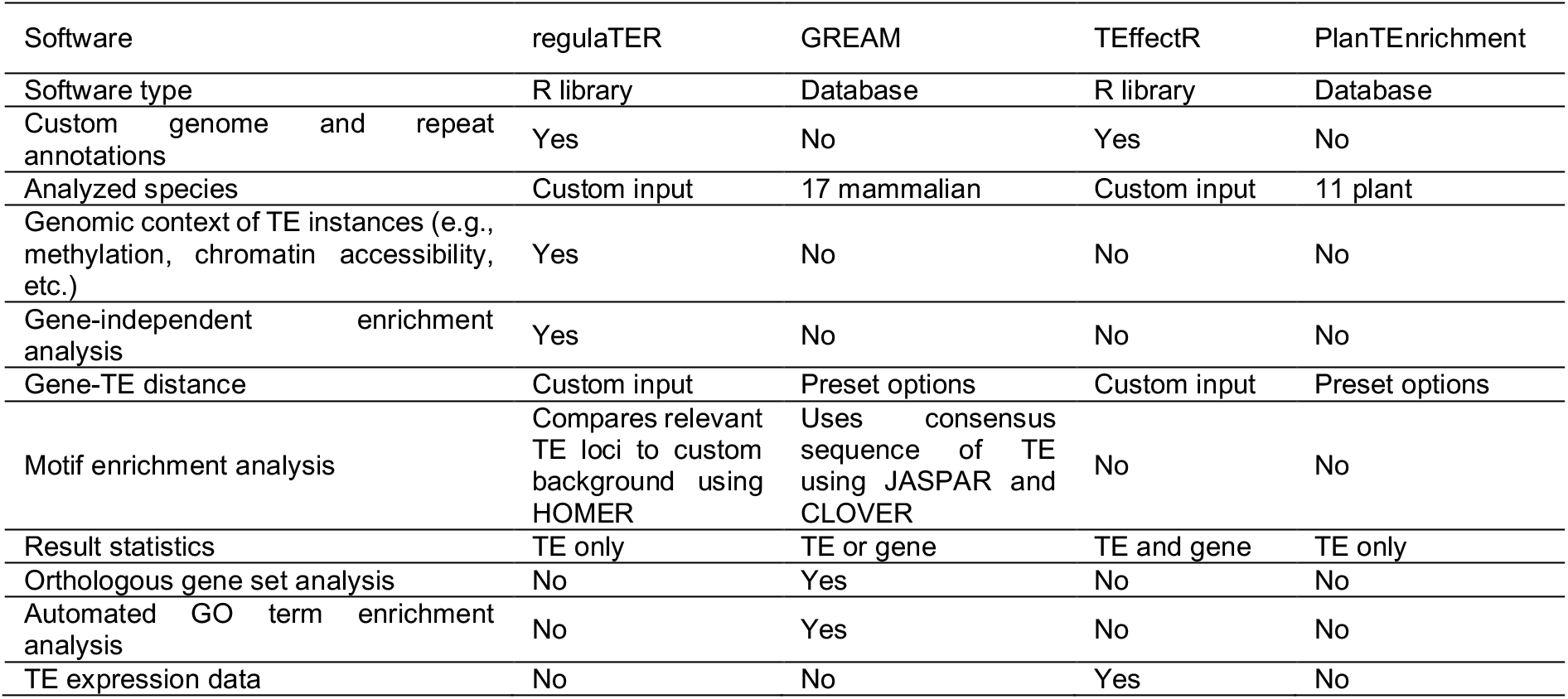
Comparison of regulaTER with similar tools.

RegulaTER can also perform TE enrichment in genomic peaks independent of gene sets using the TEAR function. Finally, regulaTER uses the HOMER software and custom background region calculation based on the multiomics input to identify motif enrichment in relevant TE loci rather than based on the consensus sequence of the TE subfamily. In comparison, GREAM only uses a limited number of preset genomes and TE annotations for its enrichment analyses. TEffectR allows for custom genome and repeat annotations but cannot include genomic peak information or perform gene-independent enrichment analysis. While the PlanTEnrichment database has the lowest functionality and output detail, it covers plant species not included in the GREAM database, which can also be used for regulaTER analyses. This comparison shows that regulaTER fills a vital niche in TE analysis tools that are not covered by other similar software.

## Discussion

In this study, we aimed to extend our understanding of the regulatory dynamics governing EMT and MET, two fundamental processes in cellular biology that play crucial roles in development, tissue repair, and disease progression, particularly cancer metastasis (Brabletz et al. 2018, Yang et al. 2020, Yao, Wang, Ma, Lin, Liu, Li and He 2024). We focused on the potential regulatory role of TEs in these processes. To analyze our EMT-MET model, we found the need to automate our analyses due to the difficulties we faced when calculating the enrichment of TEs in accessible genomic regions. Such calculations require shuffling operations that are repeated more times than is feasible to perform manually (Metropolis et al. 1953). Therefore, we developed and applied regulaTER, a unique R library meticulously crafted to dissect the regulatory roles of TEs using multi-omics data.

Previously available methods have failed to meet our analysis criteria for custom multi-omics data. For example, Chandrashekar et al. have published GREAM, a web-based tool to predict TE contributions to gene regulation by identifying their over- or under-representation near a given set of genes (Chandrashekar, Dey and Acharya 2015). However, GREAM does not consider whether the TE instances in the proximity of a gene are under a suitable chromatin state or epigenetic regulation to infer TF binding. Furthermore, it does not account for polymorphic insertions and can only perform analyses for specific reference genomes available on the web server. In addition, we have previously made available PlanTEnrichment, a tool to identify TEs enriched upstream of a list of given genes in 11 reference plant genomes (Karakulah and Suner 2017), and RTFAdb, a database of computationally predicted associations between TFBSs for 596 TFs identified using publicly available ChIP-seq data and retrotransposon species in both mouse and human genomes (Karakulah 2018). As with GREAM, however, our previous tools were limited by the number of genomes they could analyze and the genomic peak and chromatin state data they were built to handle. On the other hand, regulaTER can quickly and efficiently perform repeated calculations for any properly formatted input and any given annotated genome without requiring tedious manual interference.

Combining a well-established transcriptomic analysis and a genome-wide accessibility profile, TEAR and DATE functions of regulaTER demonstrated the enrichment of TEs in accessible genomic regions and near differentially expressed genes during EMT and MET. The FindMotifs function predicted the transcription factor bind sites within the identified TEs. Of note, major transcription factors involved in the regulation of EMT were identified, such as FoxA1/2, TEAD2, TCF/LEF, Smad3/4, Twist2, and Zeb2 (Chi et al. 2022, Gregory et al. 2008, Liu et al. 2021, Park et al. 2008, Sanchez-Tillo et al. 2011, Sayan 2014, Wang et al. 2013, Yang et al. 2004, Zavadil et al. 2004). These transcription factor binding sites were in proximity to genes that shape the mesenchymal state during EMT, such as Cdh2, Fn1, Itga5, Wnt2b, Zeb1/2, and Tgfbr1 (Gonzalez and Medici 2014, Huang et al. 2015). These findings provide unprecedented insights into the regulation of EMT through TEs, corroborating previous findings by identifying TEs enriched in accessible genomic regions during EMT; this presents a valuable validation to the capabilities of regulaTER. Several studies suggested the involvement of TEs in EMT and, thus, metastasis progression. One study used a mutagenesis approach to identify TEs involved in regulating genes driving HCC metastasis to facilitate drug identification (Kodama et al. 2016). In another study, LINE-1 elements enhanced the malignancy of human bronchial epithelial cells through interaction and co-regulation with TGFβ1 (Reyes-Reyes, Aispuro, Tavera-Garcia, Field, Moore, Ramos and Ramos 2017). Stemness and cellular plasticity during EMT in colorectal and breast cancer tumorigenesis were also shown to be regulated by the L1 retrotransposon L1-ORF1 (Apostolou, Toloudi, Chatziioannou, Kourtidou, Mimikakou, Vlachou, Chlichlia and Papasotiriou 2015). Moreover, identifying genomic insertions of AluSx1, LTR33, L2b, and MLT1F1 led to the activation of transposable element-driven transpochimeric gene transcripts such as POU5F1B, which in turn promoted colorectal cancer growth and metastasis (Simo-Riudalbas, Offner, Planet, Duc, Abrami, Dind, Coudray, Coto-Llerena, Ercan, Piscuoglio, Andersen, Bramsen and Trono 2022).

The validation of regulaTER outcomes for EMT-related TEs lays the groundwork for understanding the intricate regulatory networks governing MET, for which a similar understanding of its regulation is lagging (Alotaibi, Basilicata, Shehwana, Kosowan, Schreck, Braeutigam, Konu, Brabletz and Stemmler 2015, Werth et al. 2010, Yang, Antin, Berx, Blanpain, Brabletz, Bronner, Campbell, Cano, Casanova, Christofori, Dedhar, Derynck, Ford, Fuxe, García de Herreros, Goodall, Hadjantonakis, Huang, Kalcheim, Kalluri, Kang, Khew-Goodall, Levine, Liu, Longmore, Mani, Massagué, Mayor, McClay, Mostov, Newgreen, Nieto, Puisieux, Runyan, Savagner, Stanger, Stemmler, Takahashi, Takeichi, Theveneau, Thiery, Thompson, Weinberg, Williams, Xing, Zhou, Sheng and On behalf of the 2020). Although the TEs-dependent regulation of MET will not provide exclusive details on how it is regulated, it will add a layer of information (besides TFs and miRNAs) to this complex mechanism (Alotaibi, Basilicata, Shehwana, Kosowan, Schreck, Braeutigam, Konu, Brabletz and Stemmler 2015, Bullock et al. 2012, Cao et al. 2017, Chen and Zhao 2021, Gregory, Bert, Paterson, Barry, Tsykin, Farshid, Vadas, Khew-Goodall and Goodall 2008, Imani et al. 2017, Li et al. 2010, Sengez et al. 2019). We identified several members of the B family of TEs to be enriched near genes upregulated during MET. These elements provided binding sites for FoxA2, which has been correlated with the inhibition of EMT (Liu, Chen, Guo, Li, Zhang, Tan, Yu and Tan 2021, Zhang et al. 2015), indirectly suggesting a role in MET or the maintenance of the epithelial state. Targets of FoxA2 are also linked to EMT and MET regulation. For example, Rab17, a small GTPase of the RAS family, was shown to guard the epithelial state by inhibiting the Stat3/HIF1a/VEGF signaling pathway (Wang et al. 2020). Another target of FoxA2 is angiotensin (Agt), which can inhibit Tgfb1-induced EMT in keratinocytes (Jihu et al. 2024).

In conclusion, regulaTER offers a powerful tool for comprehensively assessing the regulatory roles of transposable elements (TEs) in gene regulation, overcoming existing limitations in the field. By applying regulaTER to multiomics data from EMT and MET processes, we identified significant contributions of the MIR and B element TE families, particularly in relation to FoxA and TEAD TFBSs near key regulatory genes. These findings underscore the nuanced regulatory roles of TEs in driving cellular transitions and phenotypic plasticity and highlight regulaTER’s ability to extend beyond traditional tools, enabling the identification of TEs’ contributions to transcriptomic regulation within specific phenotypes. This advancement represents a significant step in unraveling the complex interactions between TEs and gene regulation, offering new insights into their impact on cellular behavior.

## Supplementary Materials

The following supporting information is available online, Figure S1: GO enrichment results for differentially expressed genes near B element subfamilies of interest.; Figure S2: Expression pattern of genes enriched in the accessible ID_B1 associated biological processes.; Table S1-S8: Excel document containing Tables S1-S8.

## Declarations

### Ethics approval and consent to participate

Not applicable

### Consent for publication

Not applicable

### Availability of data and materials

RegulaTER is available at https://github.com/karakulahg/regulaTER. The ATAC-Seq and RNA-seq raw data is available at the Sequence Read Archive with the BioProject accession number PRJNA1129145.

### Authors’ contributions

D.E.: Conceptualization, Data curation, Formal analysis, Methodology, Software, Validation, Visualization, Writing—original draft, Writing—review & editing. S.Y.: Formal analysis, Investigation, Methodology, Writing—review & editing. N.A.: Conceptualization, Data curation, Formal analysis, Methodology, Software. G.K.: Conceptualization, Data curation, Formal analysis, Funding acquisition, Methodology, Project Administration, Resources, Software, Validation, Visualization, Writing—original draft, Writing—review & editing. H.A.: Conceptualization, Formal analysis, Funding acquisition, Investigation, Methodology, Project Administration, Resources, Supervision, Visualization, Writing—original draft, Writing—review & editing.

## Acknowledgments

We thank Ahmet Bursalı and Alirıza Arıbaş for their assistance with data analysis and visualization.

