## Supplementary material for "regulaTER: An R Library to Study the Regulatory Roles of Transposable Elements": Figure S1

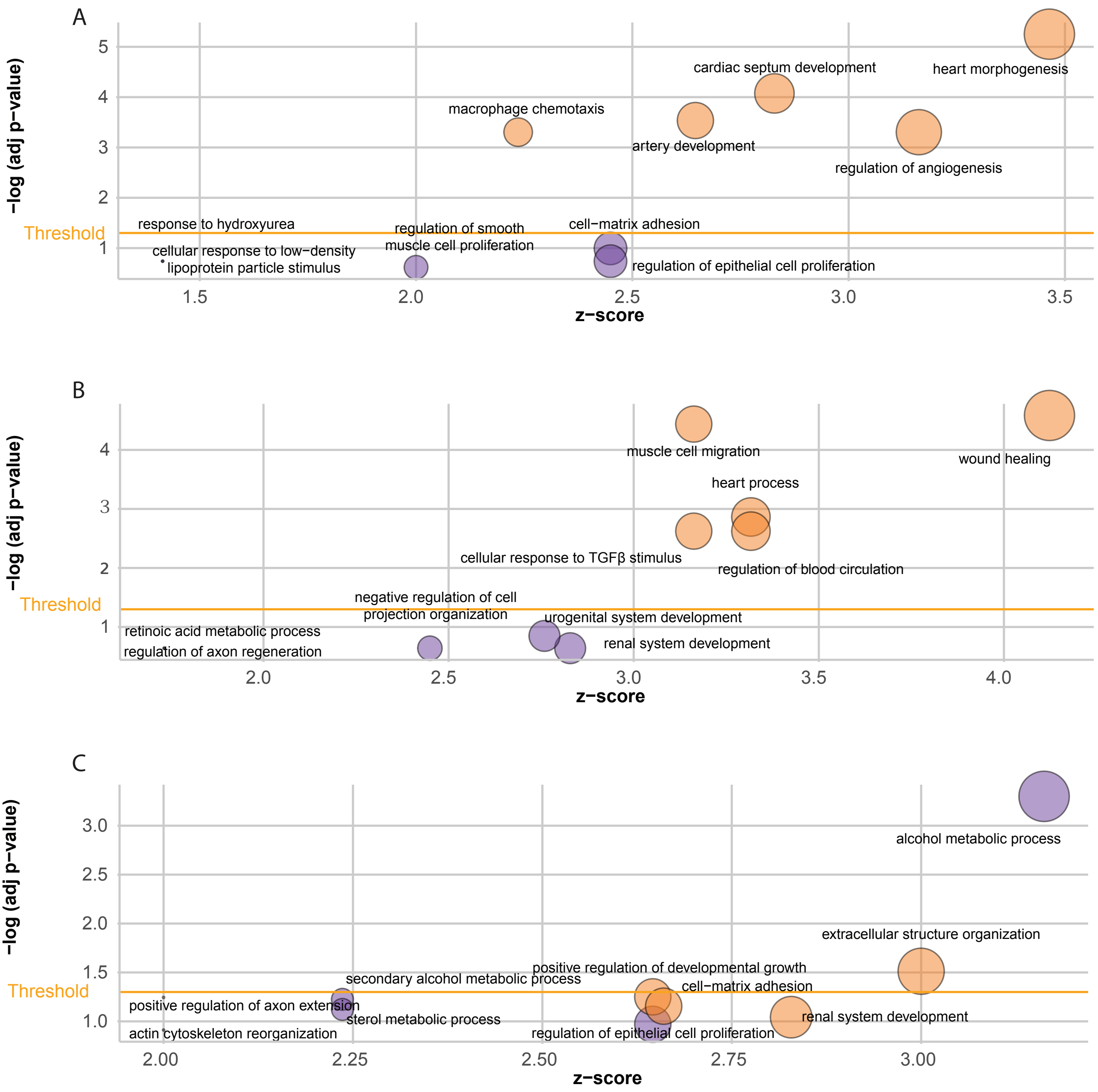


Figure S1. Five commonly enriched TEs show relevance to EMT- and MET-related biological processes. Modified GO bubble plots of the GO enrichment results for the upregulated genes within 5000 bp of the accessible instances of the B element TE subfamilies B1_Mus1 (A), B3 (B), and B3A (C). The x-axis indicates the z-score of the term, which indicates whether the genes assigned to the term are more upregulated (z-score > 0) or downregu-lated (z < 0). The y-axis represents the negative log of the adjusted p-value of the term, with the horizontal green line corresponding to an adjusted p-value of 0.05. The color of the bubbles indicates the process the term is associated with, with orange shapes enriched in EMT and purple shapes enriched in MET. The shape's size indicates the number of TE-neighboring upregulated genes in the term.
