## Supplementary material for "regulaTER: An R Library to Study the Regulatory Roles of Transposable Elements": Figure S2

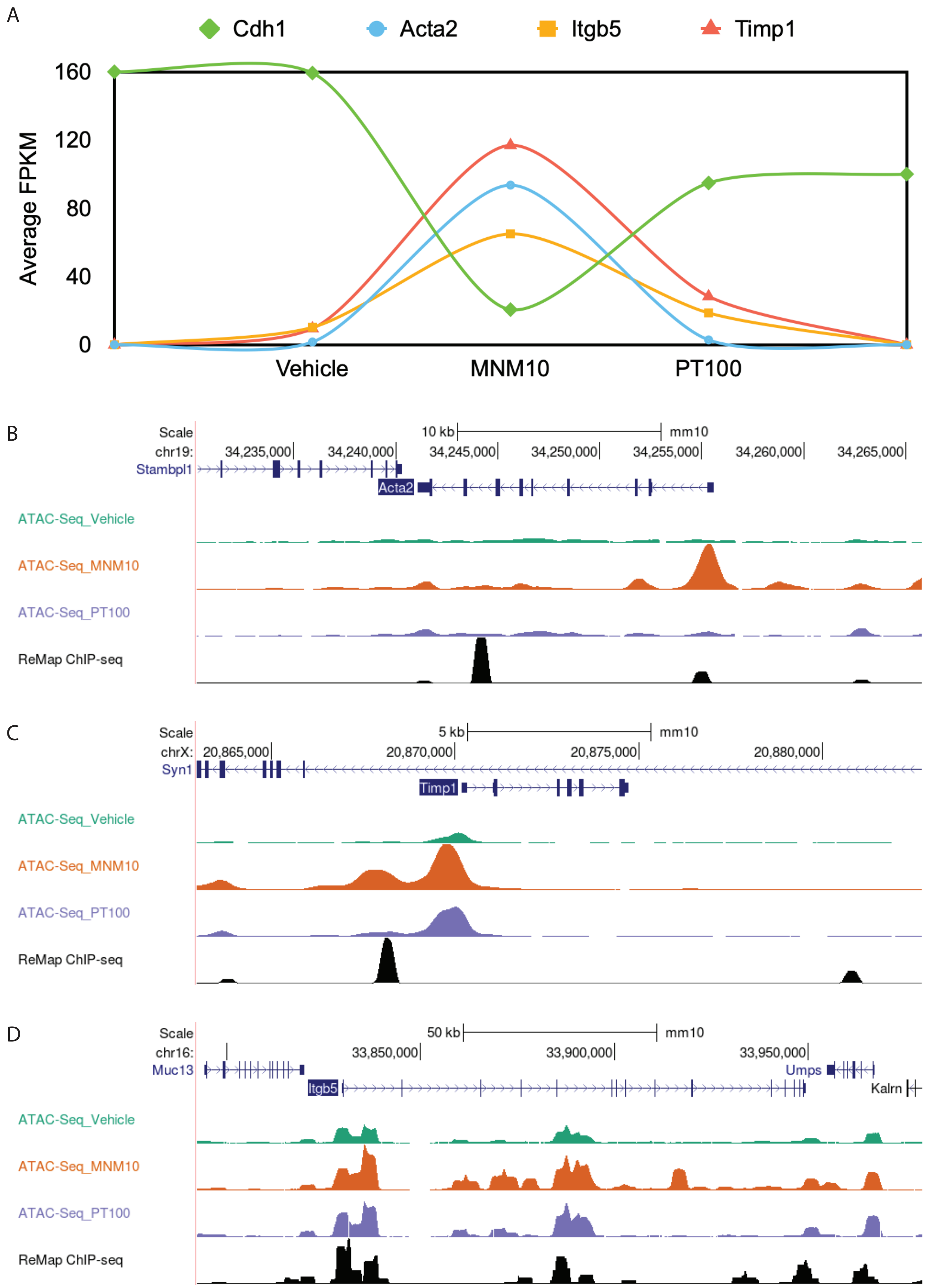


Figure S2. Expression pattern of genes enriched in the accessible ID_B1 associated biological processes. (A) Expression level changes during EMT and MET as observed in the RNA-seq data, plotted as an averaged FPKM values of the replicates. Chromatin accessibility for Acta2 (B), Timp1 (C), and Itgb5 (D) during EMT and MET as observed from ATAC-seq data and visualized in the Genome Browser. The black track indicates the ReMap ChIP-seq binding profiles for FoxA.
